# Evaluation of a Rapid Assessment Function to aid monitoring and management of Common Ravens (Corvus corax) in Washington state

**DOI:** 10.1101/2025.01.27.635125

**Authors:** Brianne E. Brussee, Shawn T. O’Neil, Michael T. Atamian, Colin G. Leingang, Peter S. Coates

## Abstract

Expanding human enterprise leading to resource subsidies for generalist species has resulted in widespread increases in common raven (*Corvus corax*) populations across the Western U.S. Ravens are an efficient predator and increased population abundance has led to adverse effects to multiple sensitive prey species. In regions where problematic interactions between ravens and their prey exist, managers seek efficient and effective tools for monitoring and controlling expanding raven populations. We previously developed a Rapid Assessment Function (RAF) for managers to quickly estimate raven population density and assess the need for management actions. We developed the RAF for the Great Basin (GB RAF) by first estimating raven density using robust distance sampling protocols with >30,000 raven point count surveys from sagebrush ecosystems in California, Nevada, Idaho, and Oregon across 131 field sites and years. We then used the relationship between raven density estimates from distance sampling and *n* ravens observed*_site-year_*/ *n* surveys*_site-year_* (that is, raven index) at each site-year combination to develop a function that accounts for detection probability and adjusts simple counts to provide a prediction of ‘true’ density. Our function produced reliable density estimates given approximately 50–100 surveys, thereby reducing the field-based and analytical efforts typically needed to estimate raven density, facilitating more efficient raven management in open sagebrush habitats. In this study, we sought to test our original GB RAF using data from sagebrush ecosystems outside of the Great Basin. Using raven point count data from two field site units in Washington state collected from 2016 to 2023, we calculated density estimates from distance sampling methods, comparable to what was done for previous analyses. We then used the GB RAF to generate predictions of density and compared those values to the more robust estimates from distance sampling. Additionally, we developed modified RAFs specifically for Washington data (WA RAFs) to assess how well they predicted raven density compared to the GB RAF. We found the detection curves estimated for the Washington sites largely aligned with those used to generate the original GB RAF. Furthermore, the estimates from the GB RAF exhibited similar or higher correlation with densities calculated from distance models (*Pearson’s r* = 0.73) than the modified WA RAFs with 1.33 km and 1.25 km truncation distances (*Pearson’s r* = 0.63 and 0.73, respectively). Producing an equivalently performing modified WA RAF would likely necessitate more data to reduce estimation error and produce more reliable estimates. These results provide evidence for the applicability of our GB RAF for more widespread use within sagebrush ecosystems, possibly negating the need for locally developed RAFs. Continued assessments of the GB RAF outside of the Great Basin would further verify its applicability across the sagebrush biome.

## INTRODUCTION

Common Ravens (*Corvus corax*; hereafter, ravens) are opportunistic omnivorous predators (Boarman and Heinrich 1999) native to North America. Across their range, raven population abundance has increased, and the areas they inhabit have expanded as increasing anthropogenic subsidies make areas accessible that were previously less inhabitable (O’Neil et al. 2018). For example, ravens have experienced elevated survival and fecundity in landscapes with human-related resource subsidies that may be used for forage opportunities or nesting structures (Kristan III et al. 2004, Boarman et al. 2006, Leu et al. 2008, Bui et al. 2010, Howe et al. 2014). Within the Cold Desert ecoregion, which includes the Great Basin and sagebrush ecosystems of eastern Washington, ravens have experienced increases in relative abundance of ∼460% from 1966 to 2018 (Harju et al. 2021).

Increases in raven abundance within the Cold Desert ecoregion can have consequences for sensitive prey species, such as the Greater sage-grouse (*Centrocercus urophasianus*; Coates et al. 2020; Coates et al. 2021), that are already challenged by human enterprise, habitat loss, changing climate and other threats (Connelly et al. 2000). Managers seeking to understand the effects of raven populations in their systems (Boarman 2003, Boarman et al. 2006, Peebles 2015) face logistical challenges regarding the survey effort needed to monitor raven abundance and track population changes through time (Dettenmaier et al. 2021). For example, 60 individual observations are recommended as the minimum number necessary to estimate a detection function based on distances of ravens from the observer (Buckland et al. 2001, Kéry and Royle 2015). A consequence of this methodology is that, especially in areas with low detection probabilities and/or densities, more surveys are needed to obtain 60 observations. Furthermore, such effort would be necessary each time a unique estimate of density was needed (for example, new management area or new year), leading to compounding effort over time.

To help alleviate the resources necessary for estimating raven density, we previously developed a rapid protocol to assess site-level density based on *n* ravens observed*_site- year_*/ *n* surveys*_site-year_* (that is, raven index; Brussee et al. 2021). We leveraged ∼30,000 raven surveys across the Great Basin encompassing 50 field sites and 131 field site-years to calculate the relationship between density estimates from robust distance sampling methods and the raven index. We then used that relationship to develop a rapid assessment function (RAF) to serve as a correction factor for detection probability on the raven index, thereby bypassing the need to conduct distance-based methodologies when logistical constraints exist. To make our RAF accessible for managers, we developed web-based software (available at https://rconnect.usgs.gov/smart/; Roth et al. 2021) that provides RAF estimates of density with uncertainty, given inputs necessary for calculating the raven index (that is, *n* ravens observed*_site-year_* and *n* surveys*_site-year_*). Based on previous findings, we posited that the RAF could generally be reliable for estimating raven density within sagebrush ecosystems characteristic of the Great Basin, given approximately 50–100 surveys (Brussee et al. 2021), which could reduce the field effort necessary to estimate raven density and facilitate more efficient raven management.

However, additional research was needed to test the predictive capacity of the RAF in sagebrush ecosystems outside the Great Basin.

To test the reliability of the Great Basin (GB) RAF outside of the region where data were originally collected, we obtained raven point count data from the Washington Department of Fish and Wildlife and the Yakima Training Center collected at two field sites within sagebrush-dominant environments in Washington state, collected from 2016 to 2023. Data have accumulated over the years and could potentially be used to estimate inter-annual variation in raven density at these regions of interest (ROIs), but research and analysis was needed to identify an appropriate, cost-effective method. Managers could benefit from a rapid but reliable method to estimate raven densities, thereby monitoring raven populations more efficiently and informing targeted and adaptive management frameworks.

Our objectives were to: 1) summarize Washington raven point count survey data to generate number of ravens and number of surveys, and for input into conventional distance sampling models; 2) evaluate the applicability of the GB RAF to predict raven densities from Washington’s ROIs (i.e., within similar environments outside of the original region of interest); and 3) use previously established methods to develop a modified RAF using only Washington data and assess the performance relative to our previously developed function. Addressing these objectives can inform the level of confidence in the accuracy of the GB RAF in ROIs in sagebrush ecosystems outside of the Great Basin and may inform future methodology to estimate raven abundance and assess the need for targeted management actions.

## METHODS

### Study area

Raven point count surveys were conducted during spring (March – June) and at two different field site units within Washington state sagebrush ecosystems (fig. 1). Both ROIs comprise a mix of shrub-steppe, rangeland, and agriculture, with relatively high potential for anthropogenic raven subsidization, likely depending on spatial proximity to point sources. The Yakima Training Center (YT) site data consisted of 149 survey locations, along a grid spaced every ∼2.3 km, where point counts were conducted once or twice per year (table 1). The Swanson Lakes Wildlife Area (SL) site unit consisted of 20 survey locations that were surveyed 7–14 times per year (table 1).

**Figure 1.**
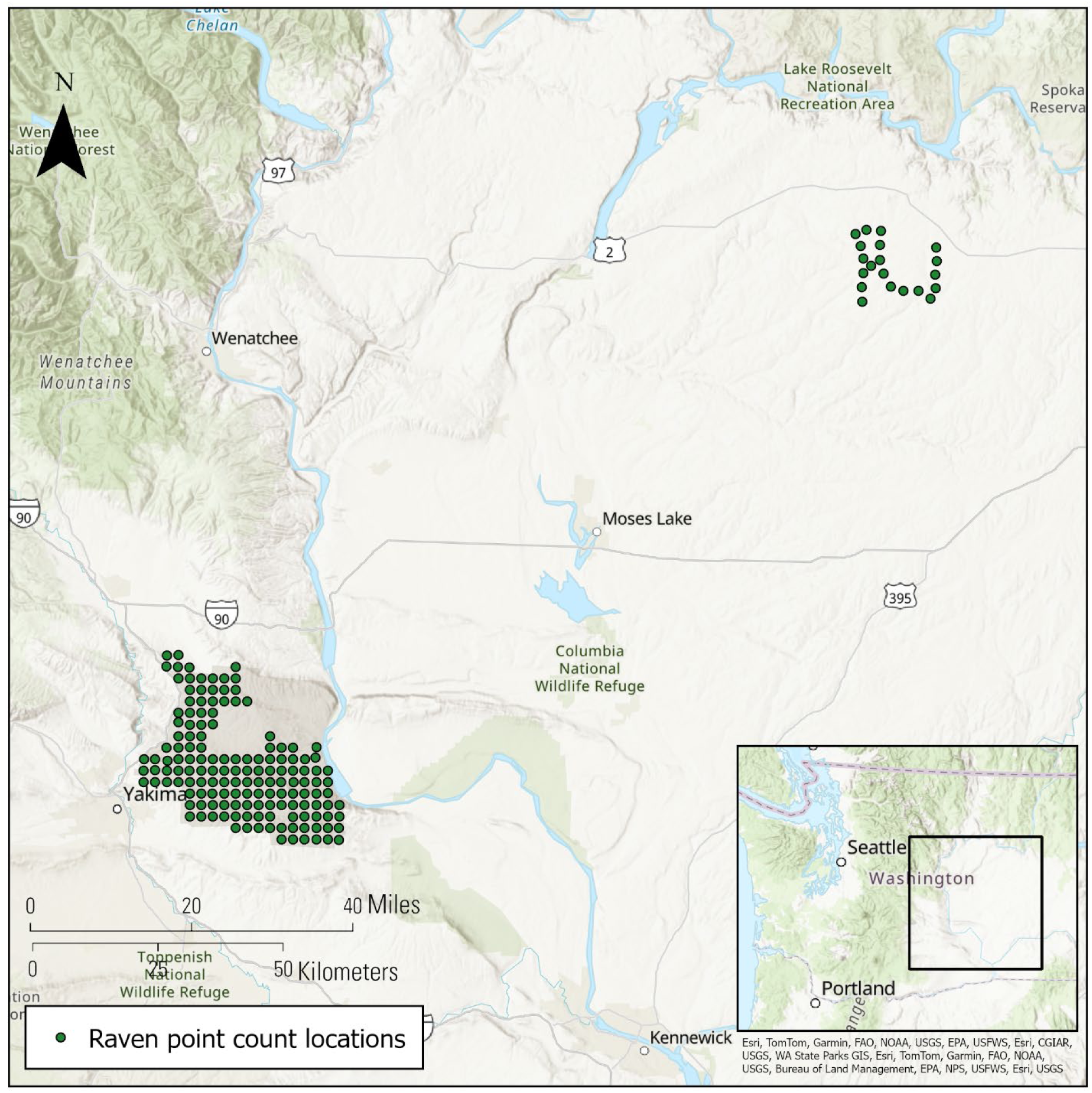
Map of raven point count survey locations at two study site units in Washington, Yakima Training Center (southwest site) and Swanson Lakes (northeast site) surveyed from 2016 to 2023.

**Table 1.** Estimates of common raven (*Corvus corax*; raven) density derived from point count data from two sites in Washington, USA from 2012 through 2023 with a truncation distance of 1.125 km. Density estimates were calculated using distance modeling approach, the Great Basin Rapid Assessment Function (GB RAF, Brussee et al. 2021), and an additional RAF calculated using a linear regression to estimate the relationship between site- and year-specific estimates of raven density from distance sampling models (ravens × km^-2^) and the average number of ravens observed per survey at the corresponding 16 site-years from Washington data. Yakima Training Center (YT), Swanson Lakes Wildlife Area (SL).

| Site | Year | N<br>ravens | N<br>surveys | N raven/<br>N survey | Distance models |  |  | GB RAF<br>(Brussee et al. 2021) |  |  | WA RAF 1.125 truncation |  |  |
| --- | --- | --- | --- | --- | --- | --- | --- | --- | --- | --- | --- | --- | --- |
|  |  |  |  |  | Est. | Lower | Upper | Est. | Lower | Upper | Est. | Lower | Upper |
| SL | 2015 | 98 | 280 | 0.35 | 0.184 | 0.141 | 0.241 | 0.304 | 0.154 | 0.599 | 0.300 | 0.112 | 0.798 |
| SL | 2016 | 127 | 280 | 0.454 | 0.215 | 0.157 | 0.294 | 0.385 | 0.195 | 0.76 | 0.429 | 0.161 | 1.144 |
| SL | 2017 | 73 | 140 | 0.521 | 0.343 | 0.247 | 0.476 | 0.438 | 0.222 | 0.864 | 0.521 | 0.196 | 1.39 |
| SL | 2018 | 91 | 220 | 0.414 | 0.38 | 0.269 | 0.538 | 0.354 | 0.179 | 0.699 | 0.378 | 0.142 | 1.007 |
| SL | 2019 | 53 | 180 | 0.294 | 0.16 | 0.114 | 0.223 | 0.259 | 0.131 | 0.512 | 0.236 | 0.088 | 0.629 |
| SL | 2020 | 25 | 63 | 0.397 | 0.244 | 0.149 | 0.4 | 0.341 | 0.173 | 0.673 | 0.357 | 0.134 | 0.95 |
| SL | 2021 | 56 | 201 | 0.279 | 0.175 | 0.127 | 0.239 | 0.247 | 0.125 | 0.486 | 0.218 | 0.082 | 0.583 |
| SL | 2022 | 98 | 155 | 0.632 | 0.467 | 0.347 | 0.629 | 0.522 | 0.265 | 1.031 | 0.682 | 0.255 | 1.821 |
| SL | 2023 | 70 | 109 | 0.642 | 0.86 | 0.58 | 1.276 | 0.53 | 0.269 | 1.046 | 0.697 | 0.261 | 1.862 |
| YT | 2012 | 79 | 147 | 0.537 | 0.879 | 0.599 | 1.29 | 0.45 | 0.228 | 0.888 | 0.544 | 0.204 | 1.45 |
| YT | 2014 | 68 | 148 | 0.459 | 0.742 | 0.507 | 1.088 | 0.39 | 0.198 | 0.769 | 0.437 | 0.164 | 1.165 |
| YT | 2015 | 82 | 126 | 0.651 | 0.948 | 0.675 | 1.331 | 0.536 | 0.272 | 1.058 | 0.71 | 0.266 | 1.897 |
| YT | 2016 | 104 | 267 | 0.39 | 0.711 | 0.55 | 0.918 | 0.335 | 0.17 | 0.661 | 0.348 | 0.13 | 0.926 |
| YT | 2021 | 32 | 148 | 0.216 | 0.235 | 0.157 | 0.353 | 0.195 | 0.099 | 0.386 | 0.153 | 0.057 | 0.412 |
| YT | 2022 | 168 | 277 | 0.606 | 0.666 | 0.511 | 0.868 | 0.503 | 0.255 | 0.992 | 0.643 | 0.241 | 1.718 |
| YT | 2023 | 80 | 286 | 0.28 | 0.275 | 0.195 | 0.387 | 0.247 | 0.125 | 0.488 | 0.219 | 0.082 | 0.586 |

### Distance model

We followed methods described in Brussee et al. (2021) using hierarchical distance sampling (Buckland et al. 2001, Thomas et al. 2010) to estimate raven density using point-count data collected at two field site units within sagebrush ecosystems in Washington. Briefly, we used the “unmarked” package (Fiske and Chandler 2011) in R 3.5.0 (R Core Team 2017) to estimate raven density for each field site and year combination using generalized distance sampling models (function ‘*gdistsamp*’) for point-count survey data (Royle et al. 2004, Sillett et al. 2012). For abundance estimates to be comparable to those used for the GB RAF, we truncated observations of ravens at 1.125 km (Coates et al. 2020, Brussee et al. 2021), and we binned distances into 5 equidistant classes (Sillett et al. 2012, Kéry and Royle 2015, Coates et al. 2020).

We specified a half-normal distance detection function to evaluate the effect of distance on detection probability (Thomas et al. 2010, Fiske and Chandler 2011). We fit area of viewshed and percent of forested covariates on the detection function, quantified as zonal means within the truncation distance (radius = 1.125 km). We modeled density using a negative binomial distribution. We specified year as covariate effects influencing abundance (Royle et al. 2004, Sillett et al. 2012) to derive densities for each year that was surveyed at each study site (Sillett et al. 2012, Kéry and Royle 2015). We assumed the distance sampling estimate of density represented (hereafter, *D*) the “true” value for the purpose of evaluating an index to approximate these densities under less rigorous sampling designs. We then calculated raven density using the raven index (*n* ravens observed*_site-year_*/ *n* surveys*_site-year_*) and the GB RAF tool to compare to the “true” densities from the distance models.

To develop a modified RAF specific to Washington data, we recalculated the truncation distance, which was where the probability of detecting ravens declined to <0.1 (Buckland et al. 2001, Burnham et al. 2004) for Washington sites (1.33 km). We then recalculated forested area and viewshed and followed all distance modeling procedures (described above) to obtain abundance estimates for the larger truncation distance.

### WA RAF development

We followed methods described in Brussee et al. (2021) for development of a RAF using only Washington data (that is, WA RAF). Briefly, for each set of results (truncation = 1.125 km and 1.33 km), we implemented a linear regression model with the log-transformed raven density estimate predicted from the natural log-transformed raven index. We report the *R^2^*and the slope coefficients from models, which were used to develop the RAFs. We compared estimates of density derived from the WA RAFs to the ‘true’ density estimates from distance models and to those from the GB RAF (1.125 km truncation distance only; Brussee et al. 2021).

Finally, we conducted an analysis to evaluate how the number of site-years used to generate a RAF affected both mean squared error (MSE) and *R^2^*of the subsequent linear regression model fits. The purpose of this analysis was to provide information on how confidence might improve with collection of data from additional site-years. To do this, we subsampled data used to develop the original RAF, and iteratively refit models. Methods and results are presented in Appendix 1.

## RESULTS

### Distance modeling results

Our analyses used data from 3,027 point count surveys conducted across two field sites in Washington sagebrush ecosystems from 2012–2023. Following Brussee et al. (2021), with a truncation distance of 1.125 km., we detected 1,304 ravens across all surveys, 691 at Swanson Lakes and 613 at Yakima Training Center. Of all surveys conducted, we detected ravens at 990 survey locations (32.7%). We detected ravens at 527 survey locations within Swanson Lakes site (32.3%) and at 463 survey locations within Yakima Training Center site (33.1%). Raven density estimated from distance sampling models ranged from 0.160–0.948 ravens per km^2^ (*x̅* = 0.463, SD = 0.284) across all field sites and years of the study (table 1). We found detection curves to approximately align with what we observed across 131 site-years from generalized distance sampling of 27 distinct regions modeled in Brussee et al. (2021) which were used to build the GB RAF (fig. 2). The raven index within each field site unit (distance ≤1.125 km) ranged 0.216– 0.651 (*x̅* = 0.445, SD = 0.142; table 1). The raven index and the distance-based model estimates of abundance were moderately correlated (*Pearson’s r* = 0.73).

**Figure 2.**
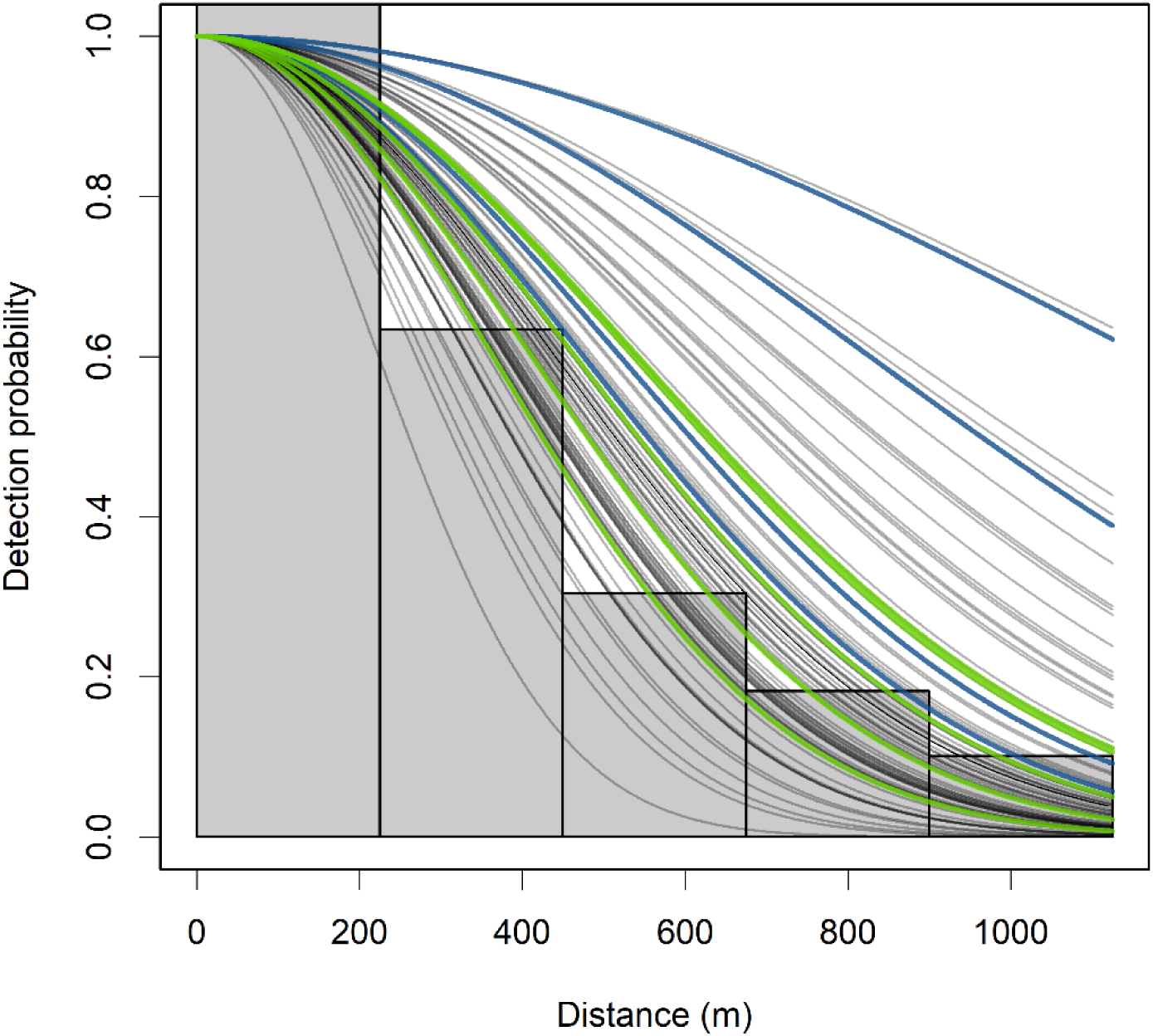
Detection curves for ravens from generalized distance sampling models (truncation distance = 1.125 km) of distinct regions across space and time. Detection curves include covariates on the detection function for area of viewshed, proportion of forested area, and year. Gray bars are frequencies of raven observations and gray lines represent each of 131 site-years from 27 regions modeled in Brussee et al. (2021). Blue and green lines are detection curves from Swanson Lakes and Yakima Training Center in Washington, USA, respectively.

When we expanded the truncation distance out to 1.33 km., we detected 1,430 ravens across all surveys, 772 at Swanson Lakes and 658 at Yakima Training Center. Of all surveys conducted, we detected ravens at 990 survey locations (32.7%). We detected ravens at 586 survey locations within Swanson Lakes site (36.0%) and at 491 survey locations within Yakima Training Center site (35.0%). Raven density estimated from distance sampling models ranged from 0.193–0.991 ravens per km^2^ (*x̅* = 0.461, SD = 0.264) across all field sites and years of the study (table 2). At the 1.33 km truncation distance, the raven index ranged 0.223–0.706 (*x̅* = 0.482, SD = 0.149; table 2). We found lower correlation between the density estimates from distance models and the raven index at the 1.33 truncation distance (*Pearson’s r* = 0.64).

**Table 2.**
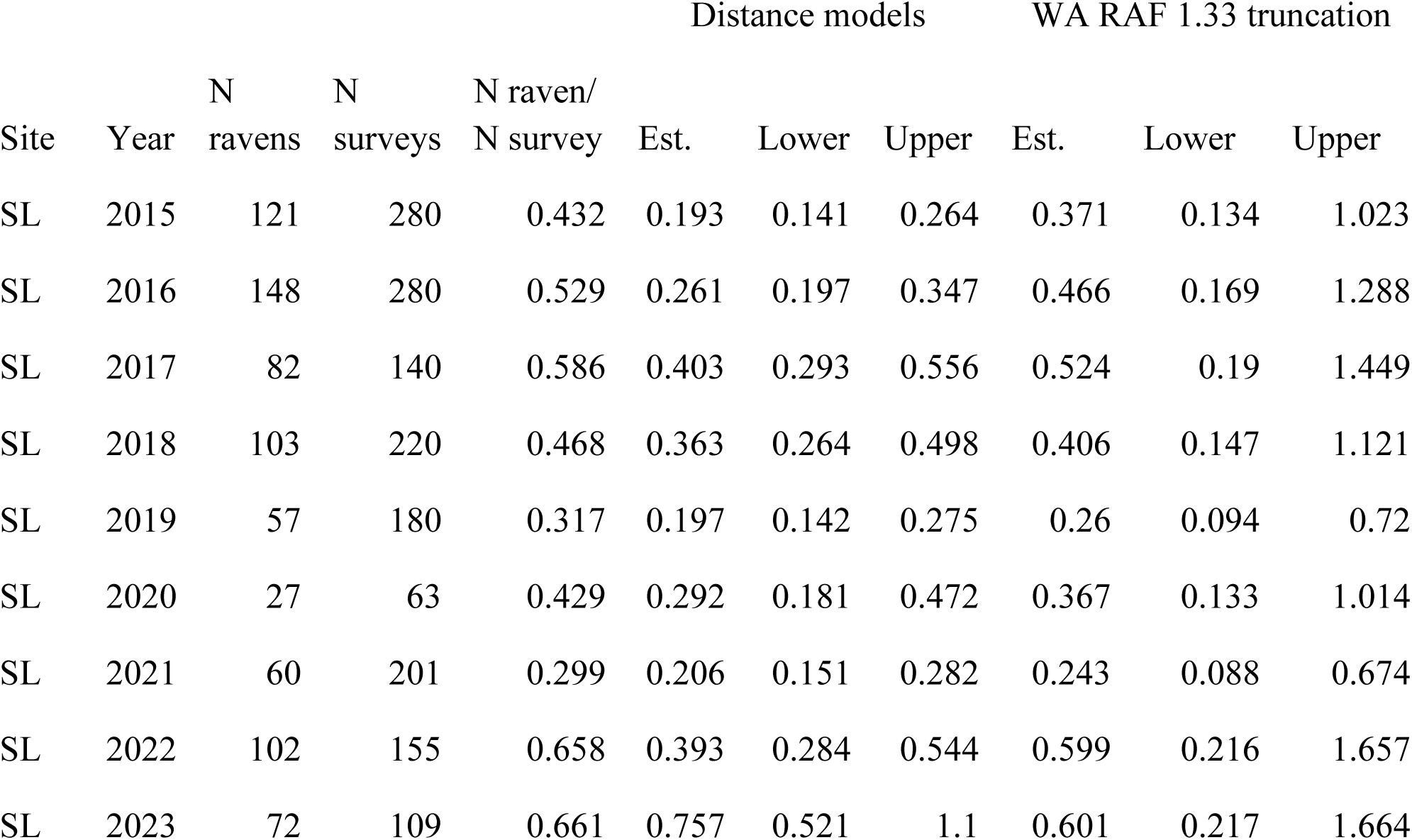
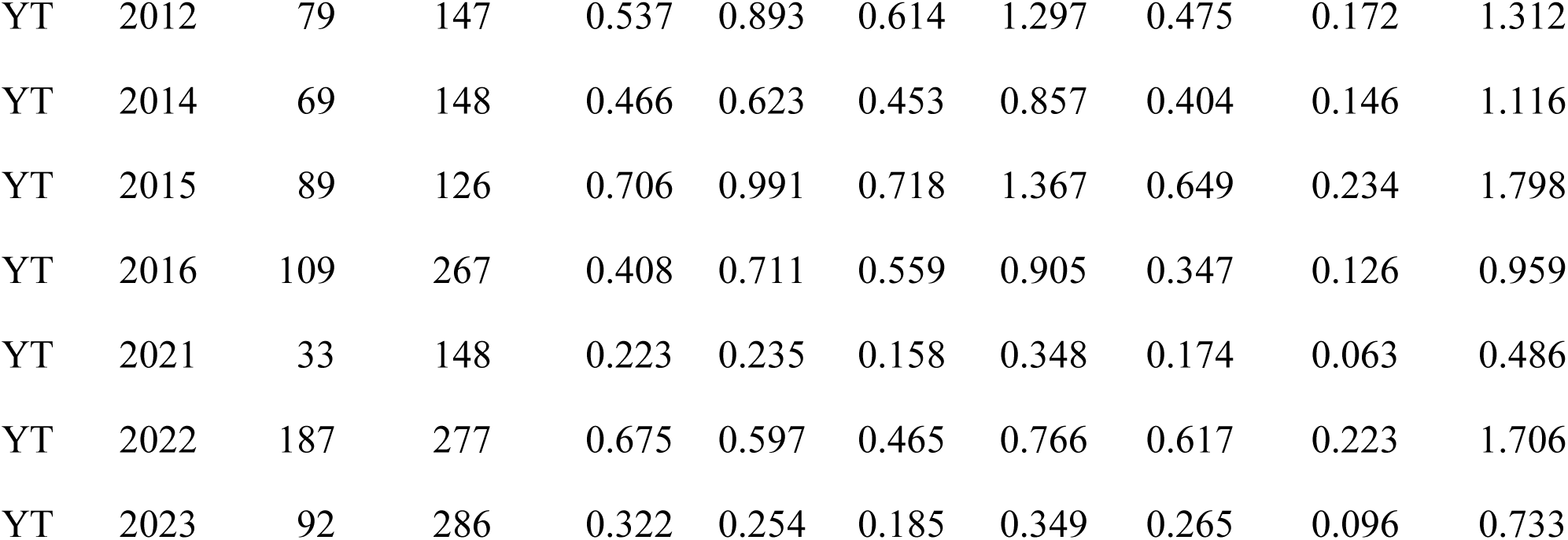
Estimates of common raven (*Corvus corax*; raven) density derived from point count data from two sites in Washington, USA from 2012 through 2023 with a truncation distance of 1.33 km. Density estimates were calculated using distance modeling approach, and a Rapid Assessment Function (RAF) calculated using a linear regression to estimate the relationship between site- and year- specific estimates of raven density from distance sampling models (ravens × km^-2^) and the average number of ravens observed per survey at the corresponding 16 site-years from Washington data. Yakima Training Center (YT), Swanson Lakes Wildlife Area (SL).

### GB RAF assessment

Brussee et al. (2021) reported a strong association between the distance-based model estimate and the raven index from the linear model (*R^2^*= 0.858) which was used to derive the GB RAF. Using the GB RAF to estimate raven density from the raven index for the Washington field site units resulted in raven density estimates ranging 0.195–0.536 (*x̅* = 0.377, SD = 0.111; table 1). We found moderate correlation between estimates from the GB RAF and those from distance models (*Pearson’s r* = 0.73; fig. 3). Correlation was stronger when assessed for individual sites, with higher correlation at the Yakima Training Center site (*Pearson’s r* = 0.88) than the Swanson Lakes site (*Pearson’s r* = 0.83). Generally, the GB RAF overestimated raven density at the Swanson Lakes site, suggesting detection was higher at Swanson Lakes than average detection observed in the Great Basin (Brussee et al. 2021; figs. 2, 3). Conversely, the RAF generally underestimated raven density at the Yakima Training Center site (fig. 3).

**Figure 3.**
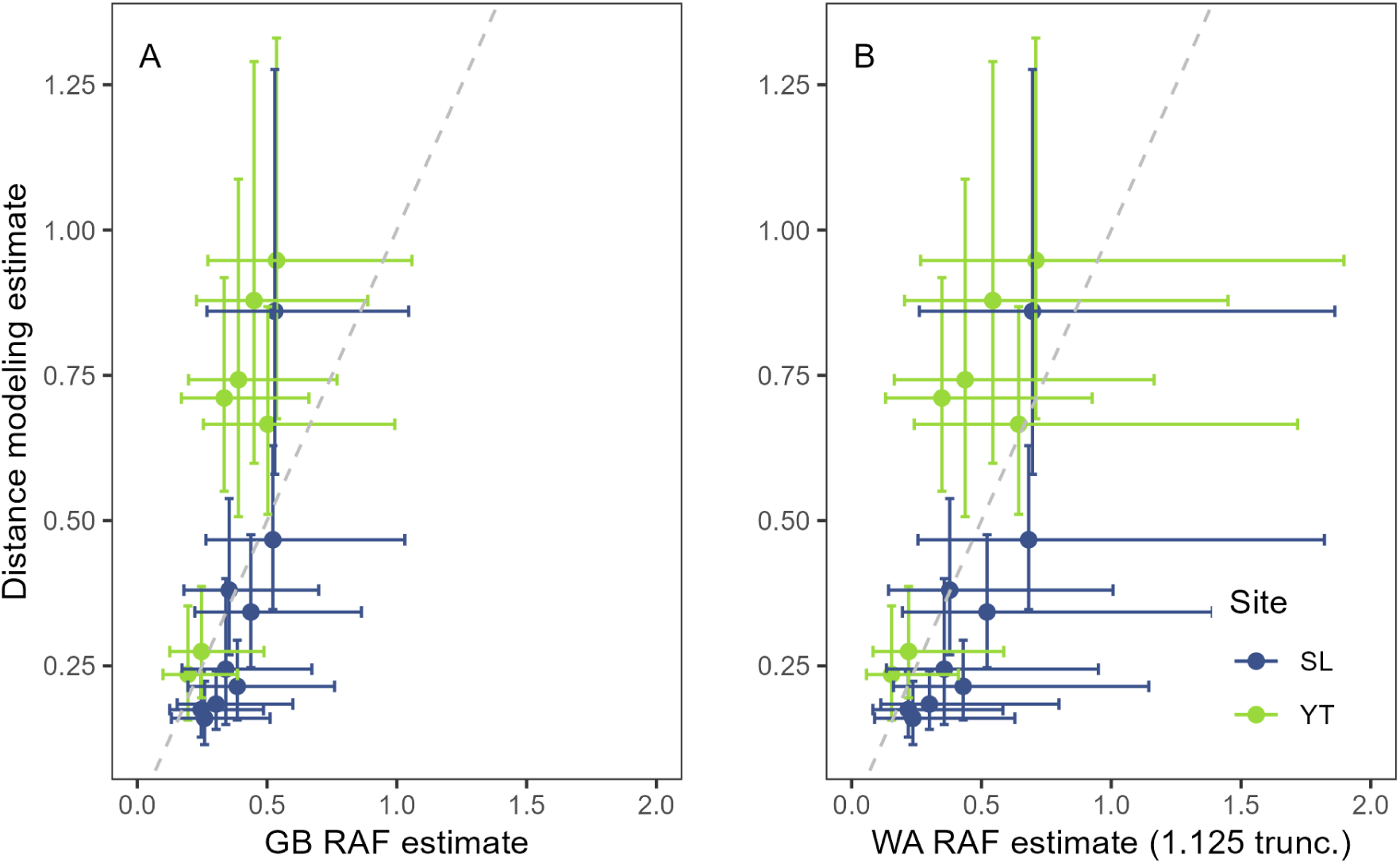
Site- and year-specific estimates of common raven (*Corvus corax*; raven) density from distance sampling models (ravens × km^-2^) with a truncation distance of 1.125 km and (A) estimates of raven density from the Great Basin Rapid Assessment Function (GB RAF; Brussee et al. 2021), and (B) estimates of raven density from a RAF derived from Washington, <u>USA</u> data at the 1.125 truncation distance. Points represent individual site-year estimates. Horizontal bars represent standard error on the RAF estimates whereas vertical bars represent standard error on the distance sampling model estimate. Dashed gray line represents a 1:1 relationship. Yakima Training Center (YT), Swanson Lakes Wildlife Area (SL).

### WA RAF development

We developed two modified RAFs for Washington data using distance sampling results with different truncation distances (figs. 4, 5; tables 1, 2). At the 1.33 truncation distance, we found the estimates from the distance models were not well approximated using the associated RAF (*β_log(ravens/survey)_* = 1.08; *R^2^* = 0.40; *Pearson’s r =* 0.63; fig. 6). At the 1.125 km truncation distance, we found the associated RAF performed similarly to the GB RAF in terms of correlation (*β_log(ravens/survey)_* = 1.39; *R^2^* = 0.52; *Pearson’s r* = 0.73; fig. 4). Given the lower sample size used to develop the WA RAFs, there was greater error associated with the abundance estimates in comparison to estimates derived from the GB RAF (fig. 3).

**Figure 4.**
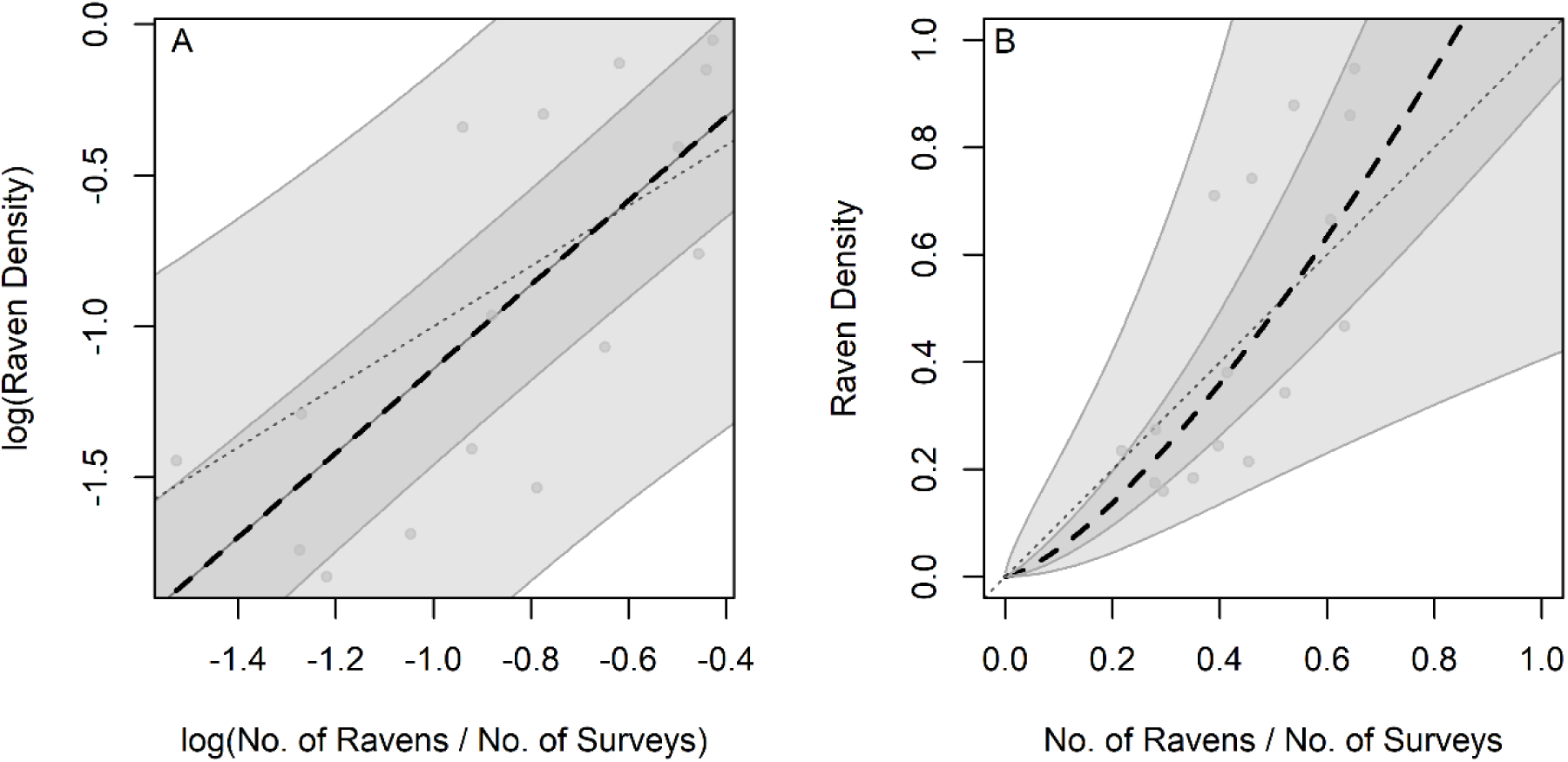
Relationship between site- and year-specific estimates of common raven (*Corvus corax*; raven) den¬sity from distance sampling models (ravens × km-2) with a truncation distance of 1.125 km and the average number of ravens observed per survey in Washington, USA at the corresponding site-years. Figure (A) represents the log-transformed data, while (B) represents back-trans¬formed estimates, which reflect output from the Rapid Assessment Function (RAF). Points represent individual site-year estimates. Light gray shading represents 95% prediction interval and dark gray shading represents 50% prediction interval. Dashed gray line represents a 1:1 relationship.

**Figure 5.**
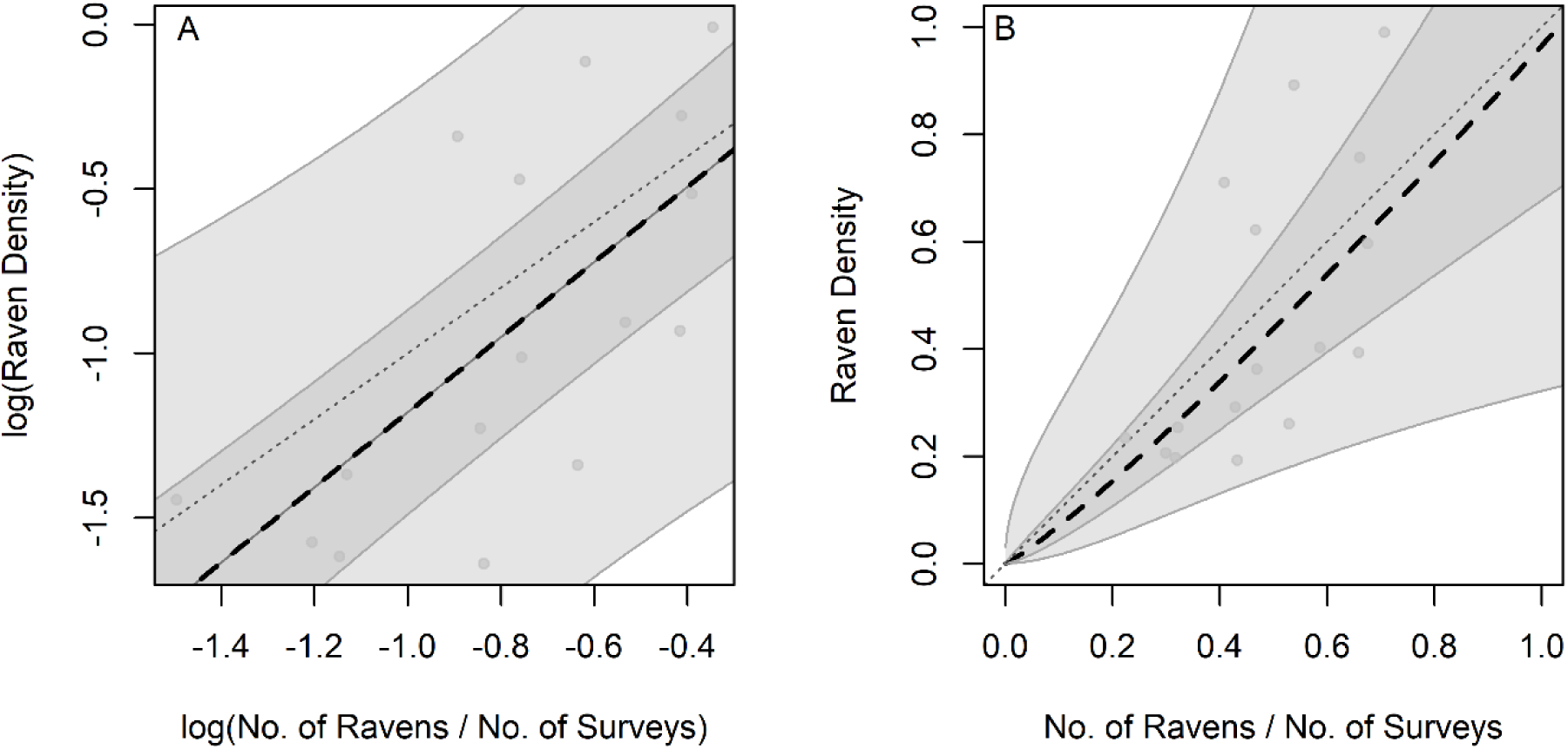
Relationship between site- and year-specific estimates of common raven (*Corvus corax*; raven) density from distance sampling models (ravens × km^-2^) with a truncation distance of 1.33 km and the average number of ravens observed per survey at the corresponding site-years in Washington, USA. Figure (A) represents the log-transformed data, while (B) represents back-transformed estimates, which reflects output from the Rapid Assessment Function (RAF). Points represent individual site-year estimates. Light gray shading represents 95% prediction interval and dark gray shading represents 50% prediction interval. Dashed gray line represents a 1:1 relationship.

**Figure 6.**
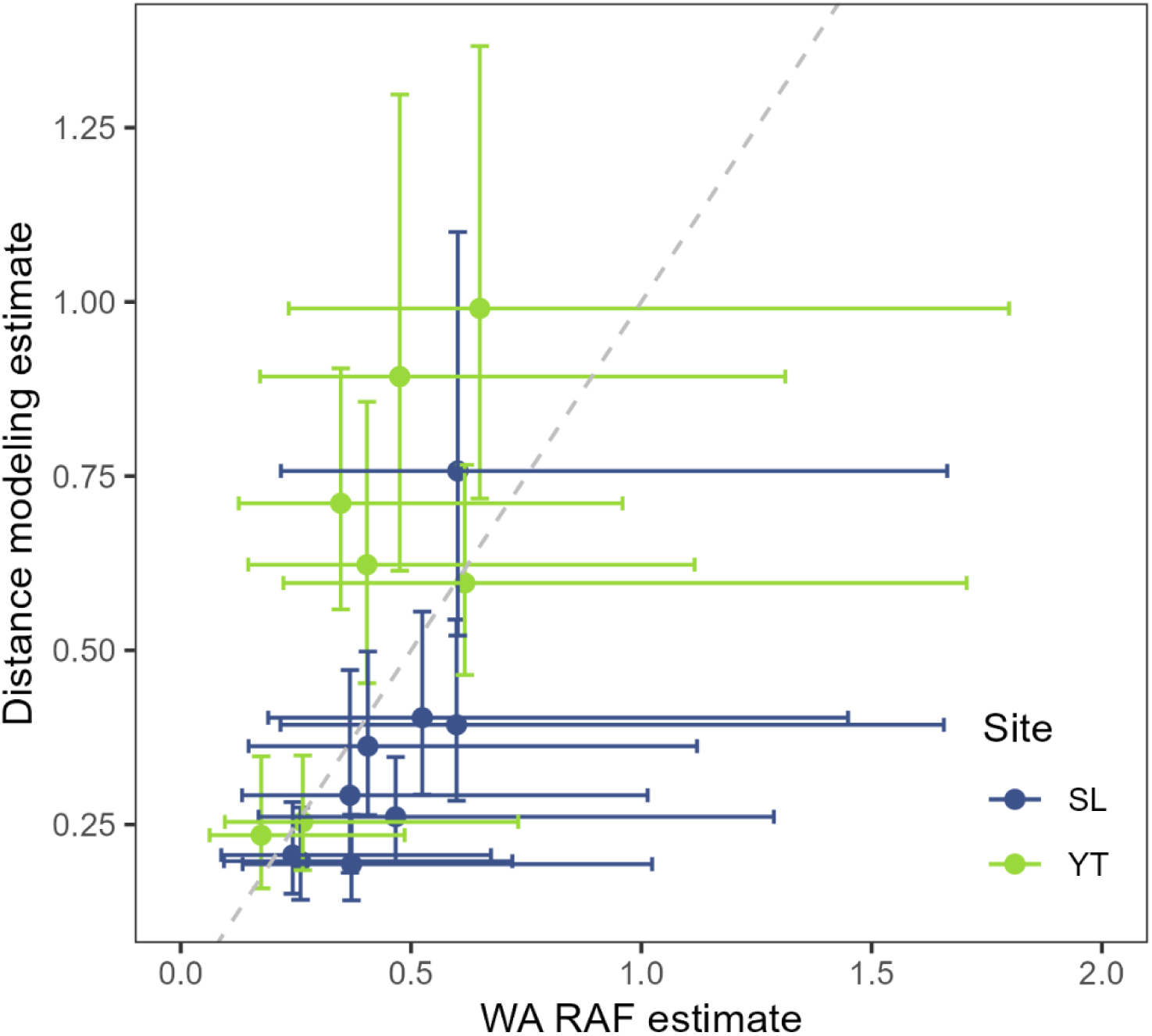
Site- and year-specific estimates of common raven (*Corvus corax*; raven) density from distance sampling models (ravens × km-2) with a truncation distance of 1.33 km and estimates of raven density from a Rapid Assessment Function (RAF) derived from Washington, USA data at the 1.33 truncation distance. Points represent individual site-year estimates. Horizontal bars represent standard error on the RAF estimates whereas vertical bars represent standard error on the distance sampling model estimate. Dashed gray line represents a 1:1 relationship. Yakima Training Center (YT), Swanson Lakes Wildlife Area (SL).

## DISCUSSION

To alleviate logistical constraints associated with conducting raven point count surveys and distance sampling analyses for assessing raven population abundance, we evaluated the applicability of our previously developed RAF within sagebrush ecosystems outside of the Great Basin. Several results from this study provide justification for use of the GB RAF in Washington sagebrush ecosystems and may inform future methodology to estimate raven abundance and assess the need for targeted management actions. First, we found the detection curves estimated for the Washington sites aligned with those used to generate the original GB RAF. Furthermore, the estimates from the GB RAF exhibited similar or higher correlation with densities calculated from distance models relative to the modified RAFs that were developed only with Washington data. The WA RAFs were developed with only 16 site-year data combinations, whereas the GB RAF leveraged ten times more point count surveys at 131 site-years. The more limited data resulted in greater error around estimates derived from the WA RAF suggesting Washington data is probably not yet robust enough for development of its own RAF. More years of data from the Washington sites could help with the development of a modified RAF, however our results provide evidence for the applicability of the GB RAF in Washington until such data are available. Alternatively, given similarities between sagebrush-dominant environments within Washington and the Great Basin, Washington data could, in the future, be combined with data from the Great Basin and elsewhere to develop a more expansive, refined RAF.

Utilizing the RAF for estimation of raven density includes the assumption of a common detection function across field sites and years and does not account for variation in detection across sites, over time, among observers, or based on sampling protocol (that is, repeat surveys or not). The Washington field sites were sagebrush steppe ecosystems characteristic of the Great Basin where the approach was developed, which are relatively open habitats where ravens are more conspicuous versus habitats characterized by dense vegetation (that is, forests). Thus, we found that the Washington sites all had detection curves that aligned with those that were used for the original RAF. However, we observed variation in detection among individual site-years which influenced how the RAF predictions approximated the distance sampling estimates. For example, compared to the detection curves observed across the Great Basin, detection probabilities were generally higher at Swanson Lakes, where a repeat sampling design was conducted. Use of a repeat sampling design, where observers visited the same locations multiple times in a single year of study, may have resulted in observers having a greater understanding of where ravens were most likely to occur at a given survey point, leading to more observances and higher detection probabilities. Therefore, the GB RAF overestimated raven at the Swanson Lakes site while it underestimated raven density at the Yakima Training Center where detection probabilities were generally lower. One caveat of density estimation from any RAF is that the function effectively pools variation in detection probabilities across space and time, thereby producing an estimate that should be interpreted as a data-informed, unbiased prediction, provided the number of new surveys is large enough (Brussee et al. 2021). As such, deviation in detection from the average detection observed across the Great Basin will affect how the GB RAF estimates density for individual site-years. However, we found it encouraging that across both Washington sites the estimates appeared to be unbiased.

For managers utilizing the GB RAF to estimate raven density in sagebrush ecosystems, other considerations include the spatial and temporal distribution of surveys. If applying the GB RAF, protocols for point count surveys should generally follow those described in Brussee et al. (2021), including the approximate size of sites, distribution of surveys within sites that include both remote areas with relatively little anthropogenic modification, and areas with more potential for anthropogenic resource subsidies. For example, the sites used to develop the GB RAF ranged in size from 42.8–4,739.1 km^2^ (average = 1,400 km^2^), and the average number of surveys per 100 km^2^ was ∼12.0 (SE = 1.7). These results suggest 50 – 100 surveys are needed to obtain density estimates for ROIs consistent in size with the sites from Brussee et al. (2021), with higher numbers of surveys conducted in larger areas to incorporate landscape variation. To best represent the variation in raven density across ROIs, randomly or systematically selected locations should include remote areas (where raven density may be relatively low) as well as areas near anthropogenic subsidies (which may be areas with higher density), with sample sites spaced to minimize potential double counting when surveys are conducted close together in time (for example, approximately 3.59 km apart, corresponding with raven home range; Smith and Murphy 1973). In addition, because of different behaviors of breeding and non-breeding ravens, timing of surveys should approximately align with the GB protocols, to span time periods of raven breeding and reproduction as raven densities may vary intra-annually. For example, reproductive ravens are largely active approximately 1,500 m around their nest, whereas when nesting concludes, large groups of ravens may congregate in areas with anthropogenic subsidies for foraging (Harju et al. 2018). Therefore, large groups of ravens may be more likely after breeding season and in areas with abundant resources. Furthermore, increases of juvenile ravens into populations following successful nesting may lead to seasonal variation in raven density estimates. While monitoring raven population trends continues to be an important component of conservation and management plans, especially in areas with increasing anthropogenic enterprise and subsequent raven densities, tools such as the RAF can advance the capabilities of land managers and biologists to monitor and respond to changing predator communities.

## ACKNOWLEDGEMENTS

Funding and logistical support for this research was provided by the Washington Department of Fish and Wildlife, with additional support from the U.S. Geological Survey Ecosystems Mission Area. Data are not currently available from the funding organization (Washington Department of Fish and Wildlife and Yakima Training Center), but can be requested by contacting Michael Atamian or Colin Leingang for more information. Any use of trade, firm, or product names is for descriptive purposes only and does not imply endorsement by the U.S. Government.

