## Appendix 1 for "Evaluation of a Rapid Assessment Function to aid monitoring and management of Common Ravens (Corvus corax) in Washington state"

This information product has been peer reviewed and approved for publication as a preprint by the U.S. Geological Survey.

**Evaluation of a Rapid Assessment Function to aid monitoring and management of Common Ravens (Corvus corax) in Washington state**

Brianne E. Brussee^1^, Shawn T. O’Neil^1^, Michael T. Atamian^2^, Colin G. Leingang^3^, and Peter S. Coates^1^

**Site-year simulation methods**

We conducted an analysis to evaluate the effect of the number of site-years on mean squared error (MSE) and *R^2^* from the linear regression models that were used to develop the RAFs. Using the log-trans­formed raven density estimates and log-transformed raven index from the Great Basin analysis, we iteratively subsampled site-year estimates with replacement, from N = 10 to N = 131, and recalculated regression models for each subset. For each sample size, we conducted 1,000 iterations, and we recorded the MSE and *R^2^*. We averaged across all iterations for each sample size to estimate how many site-years were needed for MSE to converge to a consistent estimateand for *R^2^* to approach an asymptote.

**Site-year simulation results**

Our results suggest around 50 site-year estimates are necessary for MSE to converge around the sample variance. Highest standard error around MSE occurred with site-years <50 (Figure 1A) . Additionally, while *R^2^* increased with increased number of site-years, the largest increases occurred with site-years <50 (i.e. steepest slope; Figure 1B).

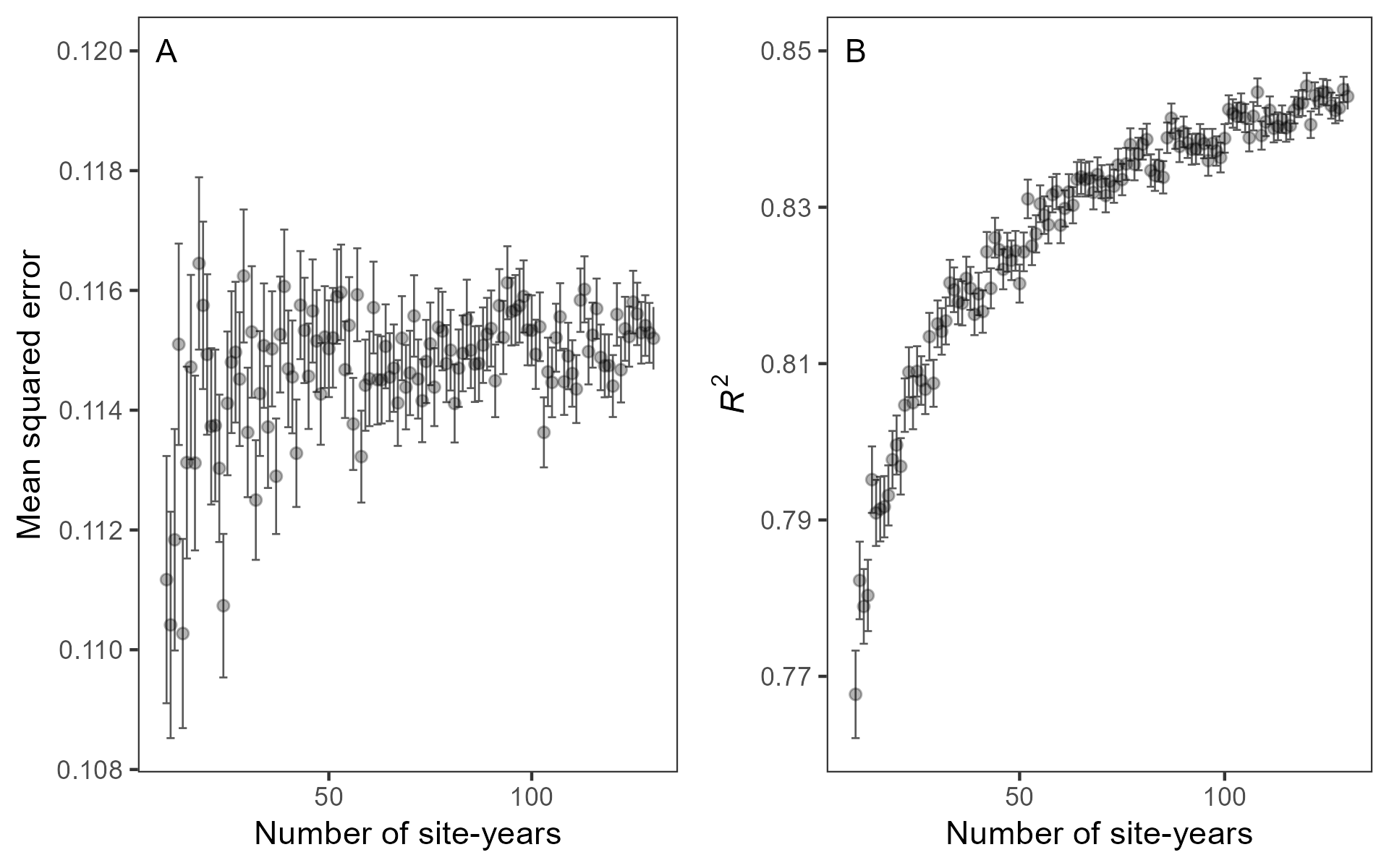

**Figure 1.** Average estimates of A) mean squared error and B) *R^2^* derived from linear regression models of raven density and the raven index from 500 iterative subsamples of site-year estimates from the Great Basin, USA (Brussee et al. 2021). Vertical bars represent the standard error.
